# Family-wide analysis of integrin structures predicted by AlphaFold2

**DOI:** 10.1101/2023.05.02.539023

**Authors:** Heng Zhang, Daniel S. Zhu, Jieqing Zhu

## Abstract

Recent advances in protein structure prediction using AlphaFold2, known for its high efficiency and accuracy, have opened new avenues for comprehensive analysis of all structures within a single protein family. In this study, we evaluated the capabilities of AphaFold2 in analyzing integrin structures. Integrins are heterodimeric cell surface receptors composed of a combination of 18 α and 8 β subunits, resulting in a family of 24 different members. Both α and β subunits consist of a large extracellular domain, a short transmembrane domain, and typically, a short cytoplasmic tail. Integrins play a pivotal role in a wide range of cellular functions by recognizing diverse ligands. Despite significant advances in integrin structural studies in recent decades, high-resolution structures have only been determined for a limited subsets of integrin members, thus limiting our understanding of the entire integrin family. Here, we first analyzed the single-chain structures of 18 α and 8 β integrins in the AlphaFold2 protein structure database. We then employed the newly developed AlphaFold2-multimer program to predict the α/β heterodimer structures of all 24 human integrins. The predicted structures show a high level of accuracy for the subdomains of both α and β subunits, offering high-resolution structure insights for all integrin heterodimers. Our comprehensive structural analysis of the entire integrin family unveils a potentially diverse range of conformations among the 24 members, providing a valuable structure database for studies related to integrin structure and function. We further discussed the potential applications and limitations of the AlphaFold2-derived integrin structures.

## 1. Introduction

Integrins are cell surface receptors that recognize a variety of extracellular or cell surface ligands, enabling communication between the cell’s interior and exterior [1]. The human integrin family consists of 24 members, formed through the combination of 18 α and 8 β subunits (**Fig. 1**). These 24 α/β integrin heterodimers are either widely distributed or specifically expressed in particular cell types. As a result, they serve universal or specialized functions in cellular processes related to cell adhesion and migration. Based on ligand or cell specificity, integrins can be categorized into subfamilies, including RGD (Arg-Gly-Asp) receptors, collagen receptors, laminin receptors, and leukocyte-specific receptors [1]. Integrins play pivotal roles in various diseases such as thrombosis, inflammation, and cancer, rendering them attractive therapeutic targets by small molecule or antibody inhibitors [2-4]. Since their discovery in the early 1980s, research into integrin structure and function has been a continuous area of interest [5].

**Figure 1.**
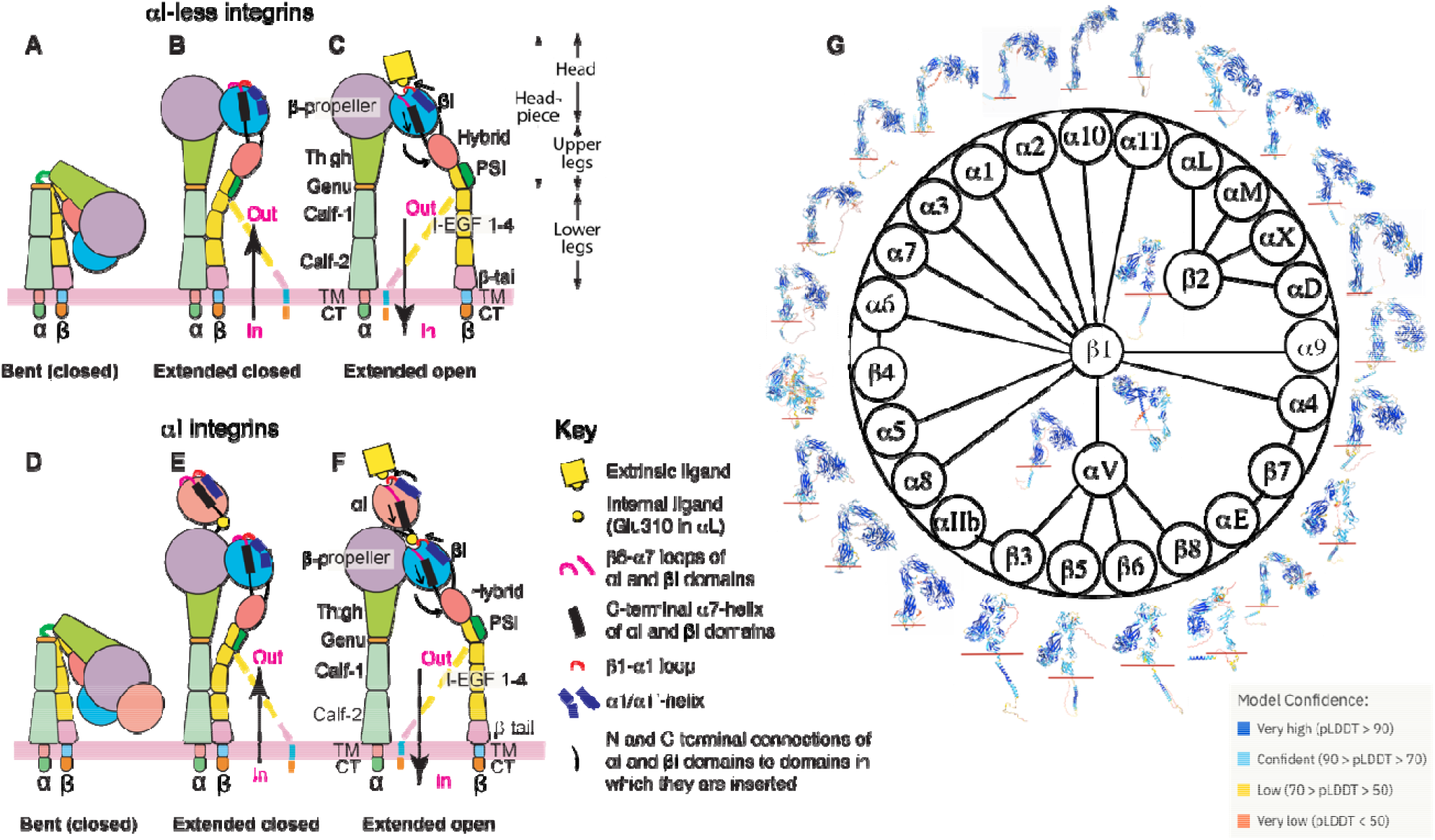
Integrin domain organization and structures predicted by AlphaFold2. (**A-F**) Integrin domain organization and conformational changes during activation are depicted for both αI-less (A-C) and αI-containing (D-F) integrins. The figures illustrate global and local structural changes, with dashed lines representing alternative conformations of the β leg domains. (**G**) The integrin family and overall structures of α and β subunits predicted by AlphaFold2 are displayed, with the red line marketing the boundary between the extracellular and transmembrane (TM) domains. The structure images are color-coded based on model confidence calculated as pLDDT scores.

The α and β subunits of integrin are composed of multiple subdomains. The α subunit contains β-propeller, thigh, calf-1, calf-2, transmembrane (TM), and cytoplasmic tail (CT) domains (**Fig. 1A-C**). The β subunit comprises βI, hybrid, PSI, I-EGF 1-4, β-tail, TM, and CT domains (**Fig. 1A-C**). A subclass of α integrins has an extra αI domain inserted into the β-propeller domain (**Fig.1D-F**). In integrins without the αI domain, the α β-propeller and β βI domains combine to form the ligand binding site (**Fig. 1A-C**). In contrast, for αI-containing integrins, the αI domain is responsible for ligand binding (**Fig. 1D-F**). Integrin ectodomains can also be divided into the headpiece (containing the head and upper legs) and lower leg domains (**Fig. 1C**). In the past few decades, structural studies of integrins have unveiled a conformation-dependent activation and ligand binding mechanism, involving transitions among at least three conformational states [6,7]. The bent conformation with closed headpiece represents the resting state of integrin (**Fig. 1A, D**), while the extended closed headpiece and extended open headpiece represent the intermediate and high-affinity active states, respectively (**Fig. 1B-C, E-F**). The conformational transition of integrin can be initiated by the binding of intracellular activators, including talin and kindlin, to β CT, leading to inside-out signaling, or by the binding of extracellular ligands, resulting in outside-in signaling (**Fig. 1A-F**) [8, 9]. However, it’s worth noting that the conformation-dependent activation model was primarily derived from structural studies of the highly-regulated β_2_ and β_3_ integrins, primarily expressed in blood cells [6,7]. Given the limited structural information available for most integrin members, it remains uncertain whether the current model of integrin conformational changes can be applied to the entire integrin family.

Since the publication of the first high-resolution crystal structure of the α_V_β_3_ ectodomain in 2001 [10], substantial efforts have been dedicated to determining integrin structures using a variety of methods, including crystallography, negative-stain electron microscopy (EM), nuclear magnetic resonance (NMR), and more recently, cryogenic EM (cryo-EM). However, high-resolution structure information remains limited to only a few integrin members, including α_V_β_3_ [10-12],^*10-12*^ α_IIb_β_3_ [13-21],^*13-21*^ α_5_β_1_ [22-24],^*22-24*^ α_6_β_1_ [25],^*25*^ α_X_β_2_ [26,27], α_L_β_2_ [28], α_M_β_2_ [29,30], α_V_β_6_ [31-33], α_V_β_8_ [34-37], and α_4_β_7_ [38], many of which have only had the fragment structures determined to date. Among the 24 integrins, the heterodimer structures of transmembrane-cytoplasmic (TM-CT) portion have been experimentally determined only for α_IIb_β_3_ [20,39-43]. The recent breakthrough in protein structure prediction using the artificial intelligence-based AlphaFold2 program has provided a powerful tool for analyzing previously challenging-to-determine protein structures with a remarkable level of accuracy [44]. We conducted an analysis of the predicted atomic structure models of single-chain 18 α and 8 β integrins that are available in the AlphaFold2 database (**Fig. 1G**). Moreover, using the recently developed AlphaFold2-multimer program [45], we predicted the structures of all 24 human integrin α/β heterodimers. Our structural analysis of the entire integrin family revealed potential conformational diversity across its 24 members, with the identification of previously unknown structural features. Our study compiled a comprehensive database of integrin structures that can serve as a valuable resource for guiding functional and structural studies. Despite the limitations of predicted structures, these findings underscore the efficacy of AlphaFold2 in the family-wide structure prediction of large and complex proteins.

## 2. Materials and methods

### 2.1. Databases and software

The single-chain structures of 18 α and 8 β integrin subunits were downloaded from the AlphaFold2 database (https://alphafold.ebi.ac.uk). Human integrin protein sequences were downloaded from the NCBI protein database (https://www.ncbi.nlm.nih.gov/protein/). Experimentally determined structures, including α_IIb_β_3_ (PDB 3FCS), α_V_β_3_ (

PDB 4G1E), α_5_β_1_ (PDB 7NXD), and α_X_β_2_ (PDB 4NEH), were downloaded from protein databank https://www.rcsb.org. All integrin structures were analyzed using PyMOL version 2.5.4 (The PyMOL Molecular Graphics System, Version 2.0 Schrödinger, LLC).

### 2.2. Running AlphaFold2

AlphaFold2 Version 2.1.2 was running on the HPC Cluster at the Medical College of Wisconsin using Miniconda3 virtual environments. The AlphaFold2 downloaded reference files are located at: “/hpc/refdata/alphafold”. Customized sbatch job script was submitted for structure prediction, shown as below.

#!/bin/bash

#SBATCH --job-name=alphafold_test

#SBATCH --ntasks=8

#SBATCH --mem=100gb

#SBATCH --time=48:00:00

#SBATCH --output=%x-%j.out

#SBATCH --gres=gpu:1

#SBATCH --partition=gpu

#SBATCH --partition=bigmem

module load alphafold/2.1.2

export NVIDIA_VISIBLE_DEVICES=$CUDA_VISIBLE_DEVICES

run_alphafold.sh -d $DOWNLOAD_DIR -f

/scratch/…/alphatest/integrin_heterodimer_amino_acid_sequence.fasta -t 2023-01-01 -o

/scratch/…/alphatest/ -m multimer

Typically, one GPU (--gres=gpu:1) and 100 GB memory (--mem=100gb) was requested to run AlphaFold2. The maximum job running time was set to 48 h (--time=48:00:00). To run AlphaFold2-Multimer for structure prediction of integrin heterodimers, an input fasta file containing the sequences of both integrin α and β subunits was provided. The multimer prediction function was enabled with command “--model_preset (-m)=multimer”. Full length or extracellular domain structures of integrin heterodimers without signal peptides were predicted with or without templates by setting the parameter of “--max_template_date(-t)=2000-05-14” or “--max_template_date(-t)=2023-01-01”. For integrin α_6_β_4_ structure prediction, the large cytoplasmic tail of β_4_ was truncated after KGRDV to simplify the prediction. The top ranked models were selected for further analysis.

### 2.3. Comparison of AlphaFold2 predicted integrin structures

The single chain α integrin structures downloaded from AlphaFold2 database were superimposed based on the α_IIb_ calf-2 domain using the “super” command in PyMOL. The single chain β integrin structures downloaded from AlphaFold2 database were superimposed based on the β_3_ βI domain using the super command in PyMOL. The experimentally determined structures for α_IIb_ (PDB 3FCS), α_V_ (PDB 4G1E), α_5_ (PDB 7NXD), and α_X_ (PDB 4NEH) were superimposed onto the predicted corresponding structures. The experimentally determined structures for β_3_ (PDB 3FCS), β_1_ (PDB 7NXD), and β_2_ (PDB 4NEH) were superimposed onto the predicted structures accordingly. The integrin heterodimer structures predicted by AlphaFold2-multimer with or without TM-CT domains were superimposed onto the calf-2 domain of α_IIb_ in PyMOL. For structure comparison of integrin TM-CT heterodimers, the structures were superimposed based on the α_IIb_ TM domain. The aligned structures were individually oriented to position them perpendicularly to the cell membrane.

### 2.4. DNA constructs

The α_5_ with C-terminal EGFP tag (α_5_-EGFP) was a gift from Rick Horwitz (Addgene plasmid #15238; http://n2t.net/addgene:15238; RRID:Addgene_15238) [46]. The α_9_ with C-terminal EGFP tag (α_9_-EGFP) was a gift from Dean Sheppard (Addgene plasmid #13600; http://n2t.net/addgene:13600; RRID:Addgene_13600). The α_3_, α_7_, and α_10_ integrins were cloned into pEGFP-N3 vector using the SalI and KpnI restriction sites following the standard molecular cloning protocol. A pair of primers were designed for introducing N839Q mutation into the α_10_-EGFP plasmid, forward primer: 5’ gaacagaaaggaaaatgcttaccagacgagcctgagtctcatcttc 3’; reverse primer: 5’ gaagatgagactcaggctcgtctggtaagcattttcctttctgttc 3’. The α_10_-N839Q mutation was generated by QuickChange mutagenesis kit following the instruction (cat# 200517-5, Agilent Technologies, Inc.). All DNA constructs were identified by sanger sequencing service provided by Retrogen, Inc.

### 2.5. Flow Cytometry Analysis of LIBS (Ligand Induced Binding Site) mAb binding

The LIBS rat mAb 9EG7 (cat# 553715, BD Biosciences) was used to measure the conformational extension of β_1_ integrin. The mouse mAb MAR4 (cat# 555442, BD Biosciences) was used to measure total surface expression of β_1_ integrin. The HEK293T cells were grown in complete DMEM (cat# 10-017-CV, Corning) supplemented with 10% fetal bovine serum (FBS) (cat# F2442, Sigma-Aldrich). Cells were maintained in a 37°C incubator with 5% CO_2_. Flow cytometry analysis of integrin expression and LIBS mAb binding were as described previously [47]. In brief, HEK293T cells were transfected with EGFP-tagged α integrin constructs plus β_1_ integrin. 48 hours post-transfection, the cells were detached, washed, and resuspended in HBSGB buffer (25 mM HEPES pH 7.4, 150 mM NaCl, 2.75 mM glucose, 0.5% BSA) containing 1 mM Ca^2+^/Mg^2+^ or 0.1 mM Ca^2+^ plus 2 mM Mn^2+^. Cells were incubated with 5 μg/ml of either 9EG7 mAb or MAR4 for 15 mins, followed by additional 15 min incubation with 10 μg/ml Alexa Fluor 647-conjugated goat anti-rat IgG (cat# A-21247, Invitrogen) or goat anti-mouse IgG (cat # A-21235, Invitrogen). Surface binding of mAb was measured by a BD Accuri™ C6 (BD Biosciences). The results were presented as a normalized mean fluorescence intensity (MFI) by calculating the MFI of 9EG7 binding (recognizing extended β_1_) as a percentage of the MFI of MAR4 binding (recognizing total β_1_). The plot was generated with Prism 9 software.

### 2.6. Data availability

The datasets generated during and/or analyzed during the current study are available from the corresponding author on reasonable request. A PyMol session file, named as “integrin-alphafold2-structure.pse”, that includes the predicted structures of integrin heterodimers were deposited online as supplementary materials.

## 3. Results

### 3.1. The AlphaFold2 single-chain structures of 18 α and 8 β human integrins

We extracted the structures of 18 human α integrins from the AlphaFold2 protein structure database. These integrin structure models in the AlphaFold2 database were predicted based on the full-length single-chain amino acid sequence, encompassing the signal peptide, ectodomain, TM and CT domains (**Fig. 1G**). The predicted models showed high accuracy in the domain structures, as indicated by the high score of predicted local distance difference test (pLDDT) (**Fig. 1G**). To compare the overall conformation of α integrin ectodomains, we superimposed all structures onto the calf-2 domain of α_IIb_, vertically oriented them relative to the cell membrane, and adjusted their rotations to display the position of the β-propeller domain relative to the membrane (**Fig. 2**). We also superimposed the AlphaFold2 structures with the experimental structures of α_IIb_, α_V_, α_5_, and α_X_. The structures were grouped based on ligand or cell specificity. Among the RGD-binding α integrins, α_IIb_ and α_V_ displayed a sharp bent conformation, nearly identical to their crystal structures (**Fig. 2A**). However, the α_5_ AlphaFold2 structure exhibited a more bent conformation than its half-bent cryo-EM structure (**Fig. 2A)**. Among the three laminin receptors, only α_6_ adopted a sharp bent conformation like the RGD receptors, while α_3_ and α_7_ showed a half-bent conformation (**Fig. 2B**). The α_4_ and α_9_ integrins also adopted a bent conformation similar to the RGD receptors (**Fig. 2C**). Interestingly, all four α integrins of collagen receptors exhibited more extended conformation than bent, with α_10_ in a nearly fully extended conformation (**Fig. 2D**). The five leukocyte-specific α integrins displayed conformational diversity, with α_L_ and α_X_ being more bent than α_M_, α_D_, and α_E_ (**Fig. 2E**). The AlphaFold2 structure of α_X_ closely resembled the α_X_ crystal structure (**Fig. 2E**).

**Figure 2.**
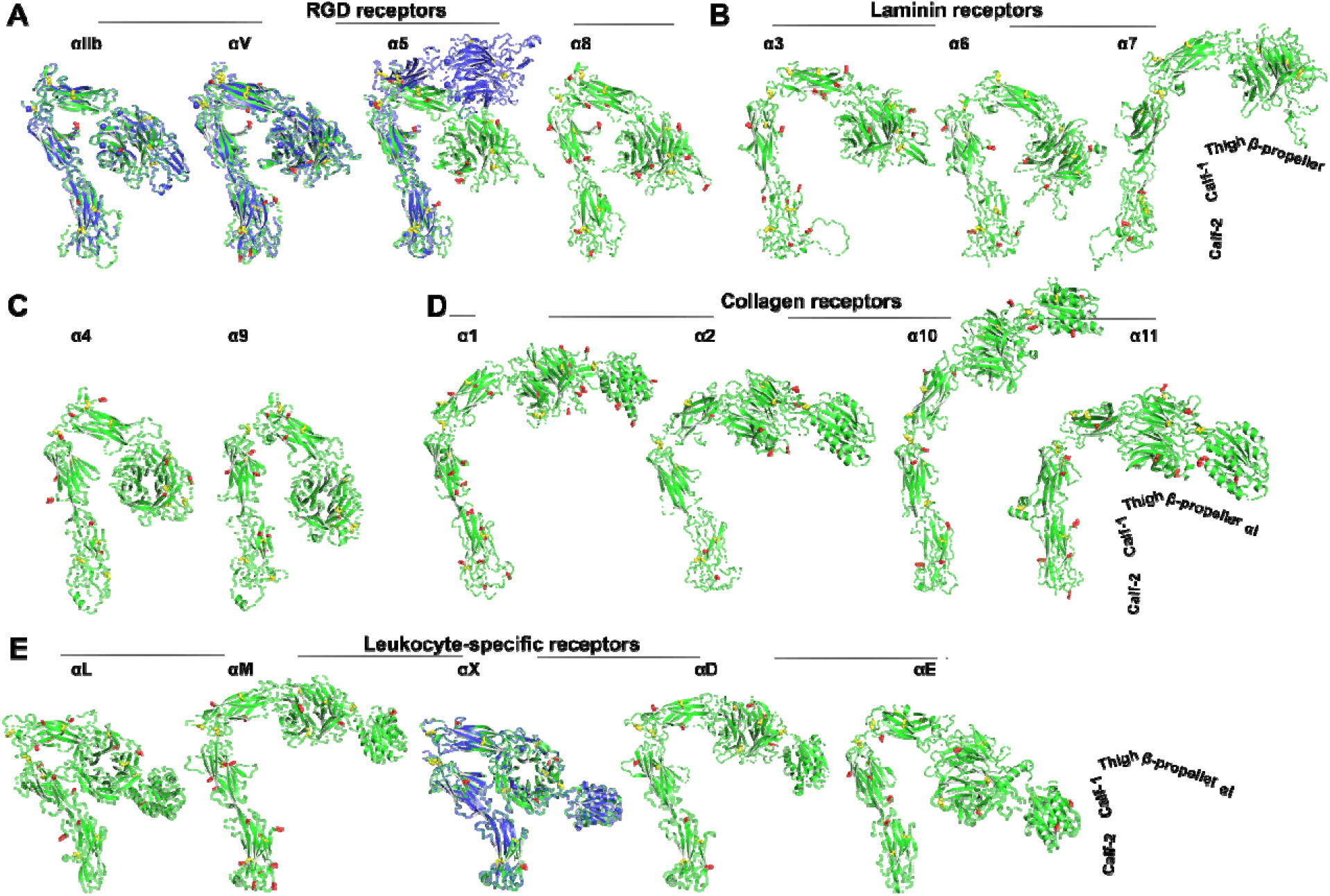
AlphaFold2 structures of the ectodomains of 18 human α integrins. The Alphafold2 structures are represented in green, while the experimentally determined structures for α_IIb_ (PDB 3FCS), α_V_ (PDB 4G1E), α_5_ (PDB 7NXD), and α_X_ (PDB 4NEH) are displayed in blue. Red sticks indicate putative N-linked glycosylation sites, and yellow sticks represent disulfide bonds. The structures were aligned based on the calf-2 domain and are oriented perpendicularly to the cell membrane.

We next analyzed the structures of 8 human β integrins, all of which were predicted as single-chain structures in the AlphaFold2 database. The predicted domain structures of β integrins also exhibit a high level of accuracy, as indicated by the pLDDT score (**Fig. 1G**). To compare the overall conformation, we superimposed the structures based on the β_3_ βI domain, and then individually orientated them vertically to membrane normal. We also superimposed the AlphaFold2 structures with the experimental structures of β_1_, β_2_ and β_3_. As depicted in **Fig. 3**, the individual domains, including βI, PSI, hybrid, I-EGF domains, and β-tail domain (β-TD), were accurately predicted for β_1_ to β_7_ integrins. However, the β-TD of β_8_ appeared smaller than other β integrins, and its structure was incompletely predicted (**Fig. 3**, β_**8**_). The AlphaFold2 structures of β_3_, β_2_, β_4_, β_5_, β_6_, and β_7_ all adopted a bent conformation as seen in the crystal structures of β_3_ and β_2_, whereas the β_1_ and β_8_ structures exhibited less bending (**Fig. 3**). The half-bent conformation of β_1_ AlphaFold2 structure closely resembled the cryo-EM structure (**Fig. 3**, β_**1**_).

**Figure 3.**
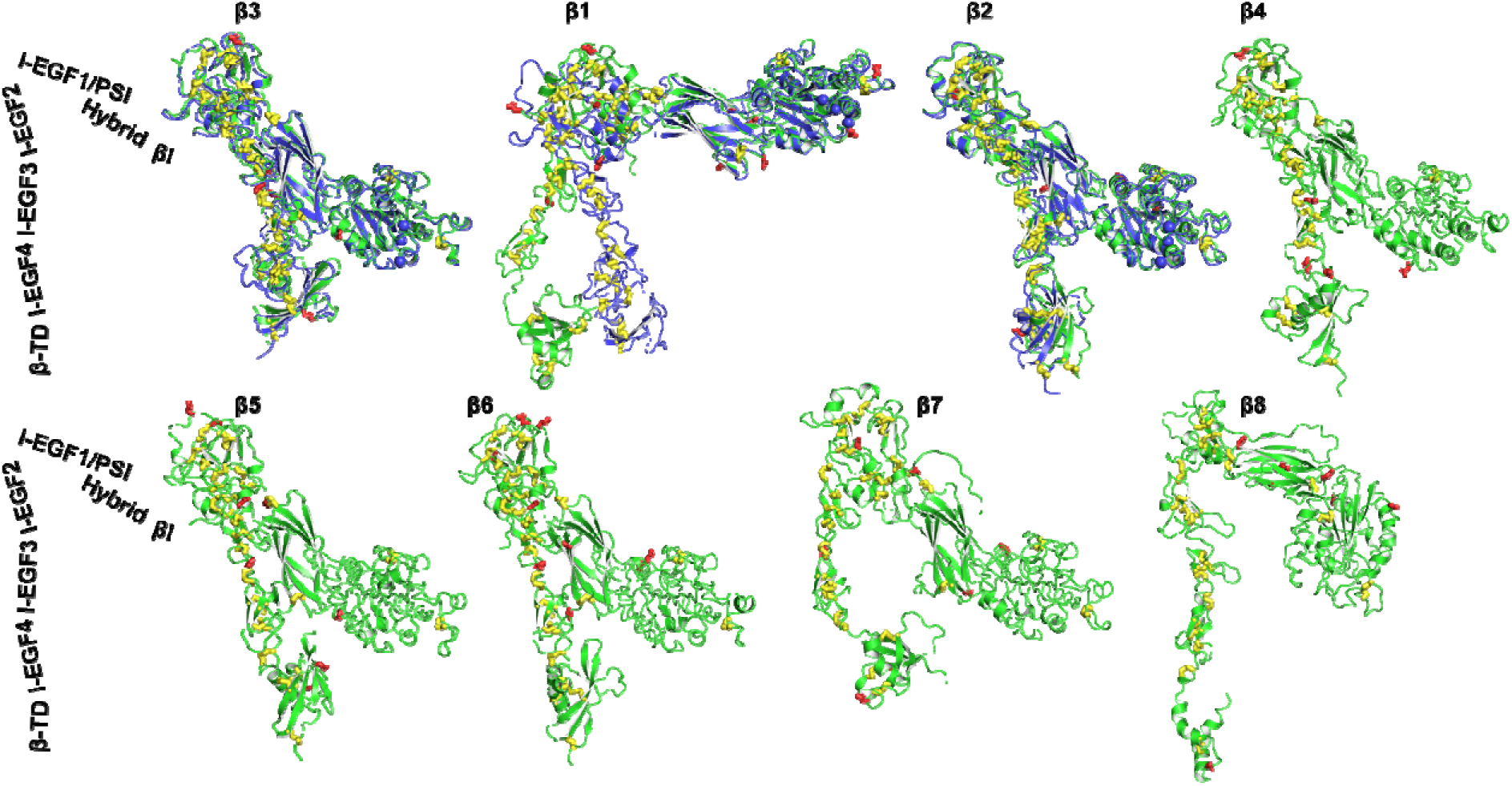
AlphaFold2 structures of 8 human β integrins. AlphaFold2 structures are depicted in green, while experimentally determined structures for β_3_ (PDB 3FCS), β_1_ (PDB 7NXD), and β_2_ (PDB 4NEH) are shown in blue. The putative N-linked glycosylation sites are represented as red sticks, and disulfide bonds are shown as yellow sticks. The structures were aligned based on the β_3_ βI domain and are oriented perpendicularly to the cell membrane.

### 3.2. The domain interface where α and β integrin subunits undergo extension

Previous structural studies have revealed that the extension of α integrin occurs at the interface between the thigh and calf-1 domains, where a disulfide bonded knob, known as the genu, is located (**Fig. 1C, F**). We conducted a sequence alignment of all α integrins at the junction between the thigh and calf-1 domains (**Fig. 4A**). To illustrate the interface between the thigh and calf-1 domains in a bent conformation, we used the structure of α_IIb_ as an example (**Fig. 4B**). Interfacial residues are highlighted in red in the sequence alignment (**Fig. 4A**) and shown as red sticks in the structure (**Fig. 4B**). The sequence alignment reveals that the interfacial residues, as well as the putative N-glycan sites, are not highly conserved (**Fig. 4A**). Some α integrins, such as α_V_, α_8_, α_4_, α_9_, α_10_, and α_E_, contain putative N-glycan sites at the interface of either thigh or calf-1 domains. Interestingly, the laminin receptors α_3_, α_6_, and α_7_ all have a longer interfacial loop (region 1) on the calf-1 domain (**Fig. 4A**). However, no signature sequences appear to indicate a preference for a bent or extended conformation.

**Figure 4.**
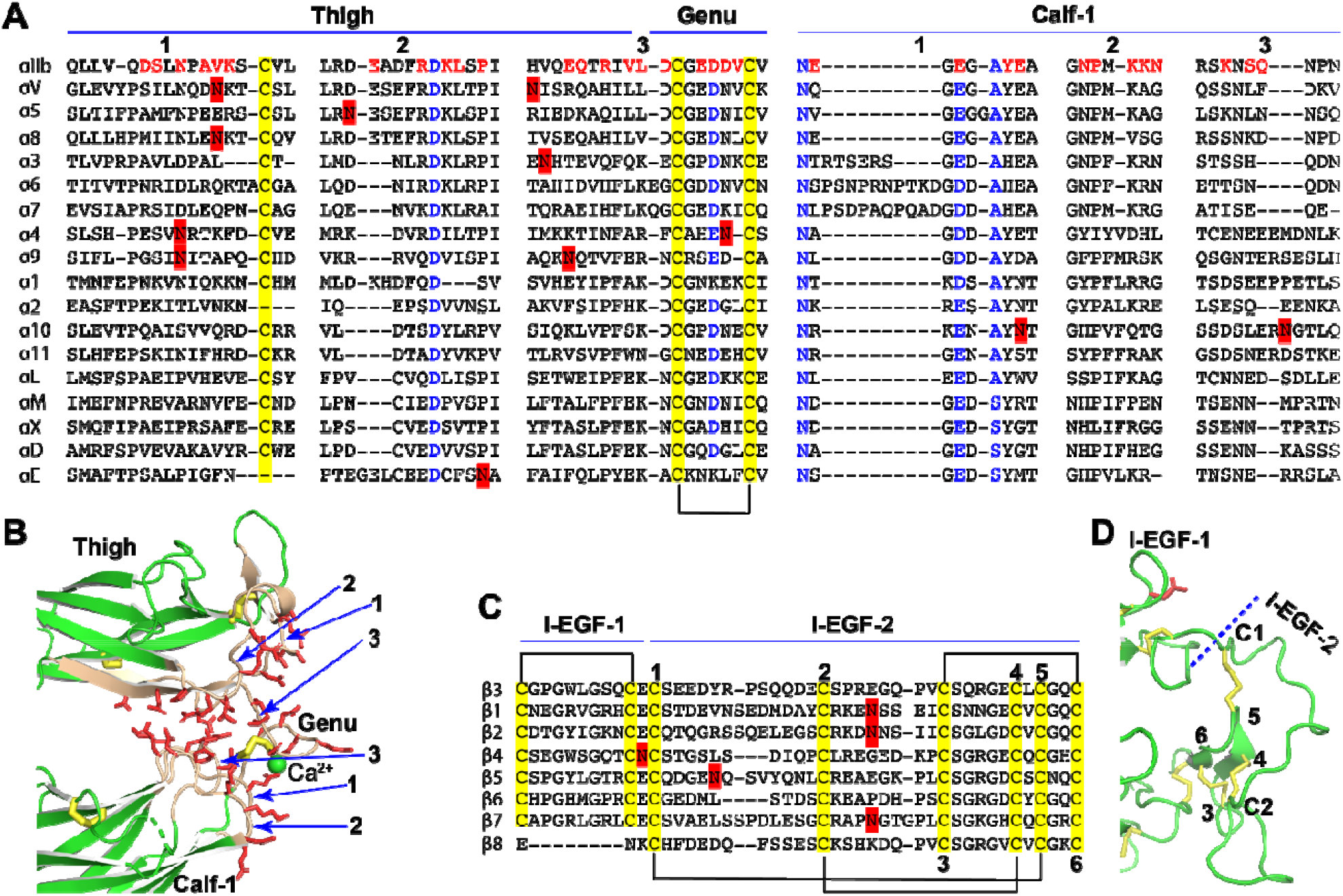
Sequence and structure of the domain interface where integrin becomes extended. (**A**) Sequence alignment of the human α integrin thigh and calf-1 domain junction interface. Interfacial residues of α_IIb_ are highlighted in red, while highly conserved residues are in blue. Disulfide bonds are indicated in yellow, and putative N-glycan sites are marked in red. (**B**) The interface between the thigh and calf-1 domains of the bent α_IIb_ structure. The loops at the interface are numbered in panels A and B. Interfacial residues are shown as red sticks. Disulfide bonds are represented by yellow sticks. (**C**) Sequence alignment of the human β integrin I-EGF-1 and I-EGF-2 domain junction. Disulfide bonds are highlighted in yellow and putative N-glycan sites are marked in red. The six cysteines of I-EGF-2 domain are numbered. (**D**) Structure of β_3_ I-EGF-1 and I-EGF-2 domain junction. Disulfide bonds are shown as yellow sticks, and one N-glycan site on I-EGF-1 is depicted as a red stick.

The integrin β subunit extends at the junction between I-EGF-1 and I-EGF-2 (**Fig. 1C, F**). Sequence alignment of the eight human β integrins in this region showed no obvious residue conservation, except for the typical disulfide bonds of EGF domains (**Fig. 4C**). In the bent conformation of β_3_ integrin (**Fig. 4D**), the interface between I-EGF-1 and I-EGF-2 is considerably smaller compared to the interface between the thigh and calf-1 in α_IIb_ (**Fig. 4B**), suggesting that it is unlikely to play a major role in maintaining the bent structure. However, the length of C1-C2 loop in I-EGF-2 domain has been shown to regulate integrin extension [48]. Notably, a landmark disulfide bond is absent in the I-EGF-1 domain of β_8_ (**Fig. 4C**), which may contribute, at least in part, to the distinct conformational regulation of β_8_ integrin.

### 3.3. α_10_ integrin prefers an extended conformation on cell surface

Among the 18 α integrins, the AlphaFold2 structure of α_10_ reveals an extended conformation (**Fig. 2D**). This extended structure was also observed in α_10_ from other species, such as mouse, rat, and zebrafish (**Fig. 5A**). Notably, we identified a conserved putative N-glycan site (N839 in human) at the interface between the thigh and calf-1 domains of α_10_ integrin (**Fig. 5A**). It is plausible that N-glycans at this site may interfere with the bent conformation (**Fig. 5B**).

**Figure 5.**
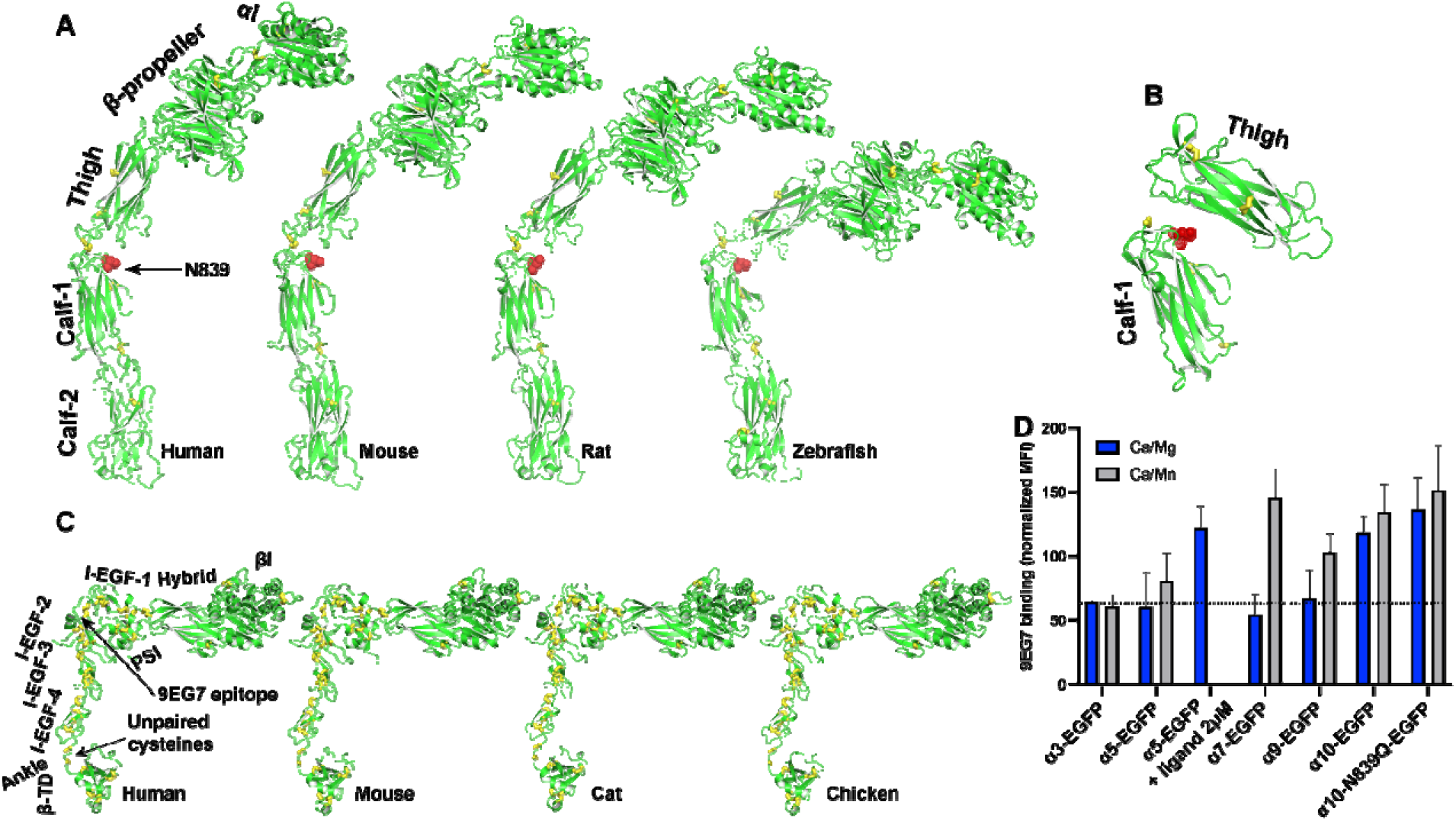
AlphaFold2 structures of α_10_ and β_1_ integrins from different species. (**A** Structures of α_10_ integrins from human, mouse, rat, and zebrafish. The structures were aligned based on the calf-2 domain and oriented perpendicularly to the cell membrane. The N-glycan site at the thigh/calf-1 interface is shown as a red stick. (**B**) Relative positions of α_10_ thigh and calf-1 domains in the fully bent conformation, illustrating the interfacial location of the N-glycan site shown as red sticks. The thigh and calf-1 domains of α_10_ were superimposed on those of bent α_IIb_ integrin structure. (**C**) Structures of β_1_ integrins from human, mouse, cat, and chicken. The structures were aligned based on the βI domain and oriented perpendicularly to the cell membrane. (**D**) Conformation of β_1_ integrin co-expressed with selected integrin α subunit. Human integrin α subunits with a C-terminal EGFP tag were co-expressed with human β_1_ in 293T cells. The binding of mAb 9EG7 or MAR4 was measured by flow cytometry in a buffer containing 1 mM Ca^2+^/Mg^2+^ or 0.1 mM Ca^2+^ plus 2 mM Mn^2+^. The data are presented as the MFI of 9EG7 binding as a percentage of the MFI of MAR4 binding.

It is important to note that α_10_ integrin only forms heterodimer with β_1_ integrin. The AlphaFold2 structures of human, mouse, cat, and chicken β_1_ integrins all exhibit a half-bent conformation (**Fig. 5C**). To measure the conformation of α_10_β_1_ on cell surface, we used mAb 9EG7, which recognizes the β_1_ I-EGF-2 epitope that remains concealed in the bent conformation (**Fig. 5C**) [49]. This antibody reports on β_1_ integrin extension. As a positive control, we used the universal integrin activator Mn^2+^ to induce integrin extension. For comparison with α_10_, we selected α_3_, α_5_, α_7_, and α_9_ integrins.

In our experiments, α integrins were expressed as EGFP-fusion proteins alongside β_1_ integrin in HEK293T cells. Flow cytometry analysis under physiological metal ion condition (1 mM Ca^2+^/Mg^2+^) revealed that α_10_-EGFP/β_1_ cells exhibited higher binding to 9EG7 compared to other integrins (**Fig. 5D**). Conversely, α_7_-EGFP/β_1_ and α_9_-EGFP/β_1_ cells showed increased 9EG7 binding only under the activating condition (0.1 mM Ca^2+^/ 2 mM Mn^2+^). In contrast, α_3_-EGFP/β_1_ cells did not response to Mn^2+^ (**Fig. 5D**). α_5_-EGFP/β_1_ cells also exhibited limited responsiveness to Mn^2+^ for 9EG7 binding compared to α_7_ and α_9_. However, the RGD-like compound MK-0429 effectively induced 9EG7 binding to α_5_-EGFP/β_1_ cells (**Fig. 5D**). Notably, Mn^2+^ did not further increase 9EG7 binding to α_10_-EGFP/β_1_ cells (**Fig. 5D**). Mutating the putative N-glycan site at the interface between the thigh and calf-1 domains of α_10_ integrin (α_10_-N839Q) did not affect 9EG7 binding (**Fig. 5D**). These data suggest that α_10_ integrin maintains a constitutively extended conformation on cell surface.

### 3.4. The integrin α/β heterodimer structures predicted by AlphaFold2-multimer

Integrins are typically expressed as heterodimers on cell surface, consisting of both α and β subunits. However, the integrin structures in AlphaFold2 database were modeled individually for the single-chain α and β subunits, as described above. To construct integrin structures containing both α and β subunits, we utilized the AlphaFold2-multimer module for structure prediction of all 24 human integrin heterodimers. To prevent potential model bias arising from the templates of experimental integrin structures, we set the template search date to the year 2000, predating the reporting of any integrin heterodimer structures. Impressively, AlphaFold2-multimer successfully predicted the structures of all 24 integrin heterodimers (**Fig. 6**). These structures were categorized based on their ligand or cell specificity. To facilitate conformational comparisons, all structures were superimposed onto the α_IIb_ calf-2 domain and individually orientated to align their ectodomains vertically with respect to the cell membrane (**Fig. 6**). Overall, the inter-subunit interfaces, including those of the α β-propeller and β βI domains, were accurately modeled. Notably, the AlphaFold2 structures of α_IIb_β_3_ and α_V_β_3_ ectodomains closely resembled their respective crystal structures, with a Cα root-mean-square deviation (RMSD) of approximately 2 Å, highlighting the outstanding capability of AlphaFold2 in modeling integrin structures. Among the RGD receptors, all adopted a bent conformation, except α_5_β_1_, which exhibited a half-bent (**Fig. 6A**). Similarly, all laminin receptors, including α_7_β_1_, displayed a bent conformation (**Fig. 6B**). Interestingly, the four collagen receptors, including α_10_β_1_, exhibited a half-bent structure (**Fig. 6C**), in contrast to the AlphaFold2 structure of single-chain α_10_ showing a more extended conformation (**Fig. 2D**). Within the leukocyte-specific integrins, only α_L_β_2_ and α_E_β_7_ displayed a sharp bent conformation, while α_M_β_2_, α_X_β_2_, α_D_β_2_, and α_E_β_7_ appeared to be more extended (**Fig. 6D**). The AlphaFold2 structures of α_4_β_1_ and α_9_β_1_ showed a half-bent conformation (**Fig. 6E**). It is worth noting that the relative orientation of the TM domains to the cell membrane was not correctly predicted for most of the structures.

**Figure 6.**
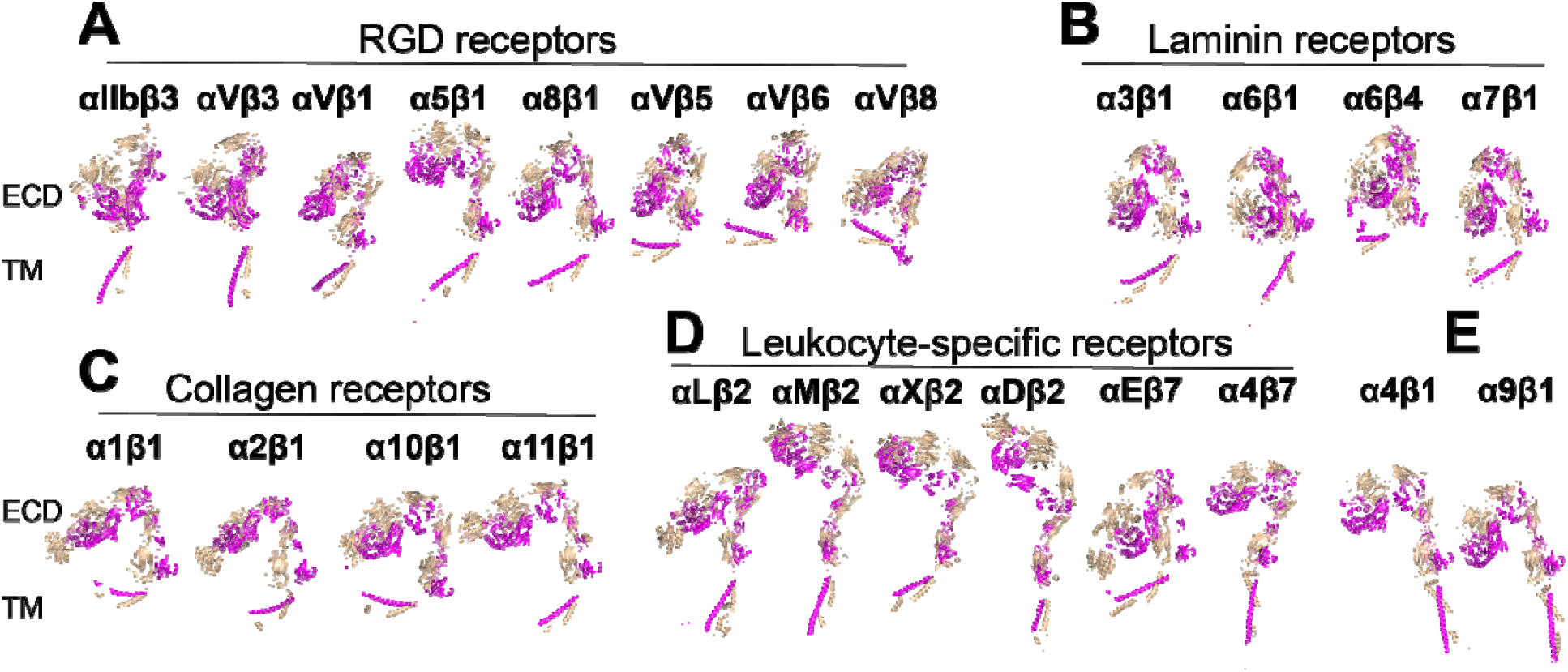
Structures of 24 human integrins predicted by AlphaFold2-multimer. (**A-E**) Full-length integrin structures were calculated using AlphaFold2-multimer with the max_template_date set to year 2000. Structures were superimposed onto the calf-2 domain of the α_IIb_ subunit and are presented in the same orientation. The α and β subunits are shown in wheat and magenta, respectively.

As nine of the integrin structures predicted by AlphaFold2-multimer show artificial interactions between TM-CT and ectodomains (**Fig. 7A**), we asked whether such interactions have any impact on the overall integrin structure prediction. We performed AlphaFold2-multimer modeling of the nine integrin structures in the absence of the TM-CT sequences. Surprisingly, the resulting structures closely matched those containing TM-CT domains (**Fig. 7B**), indicating that the structure modeling of the ectodomain and TM-CT domains does not influence each other during the structure calculation by AlphaFold2-multimer.

**Figure 7.**
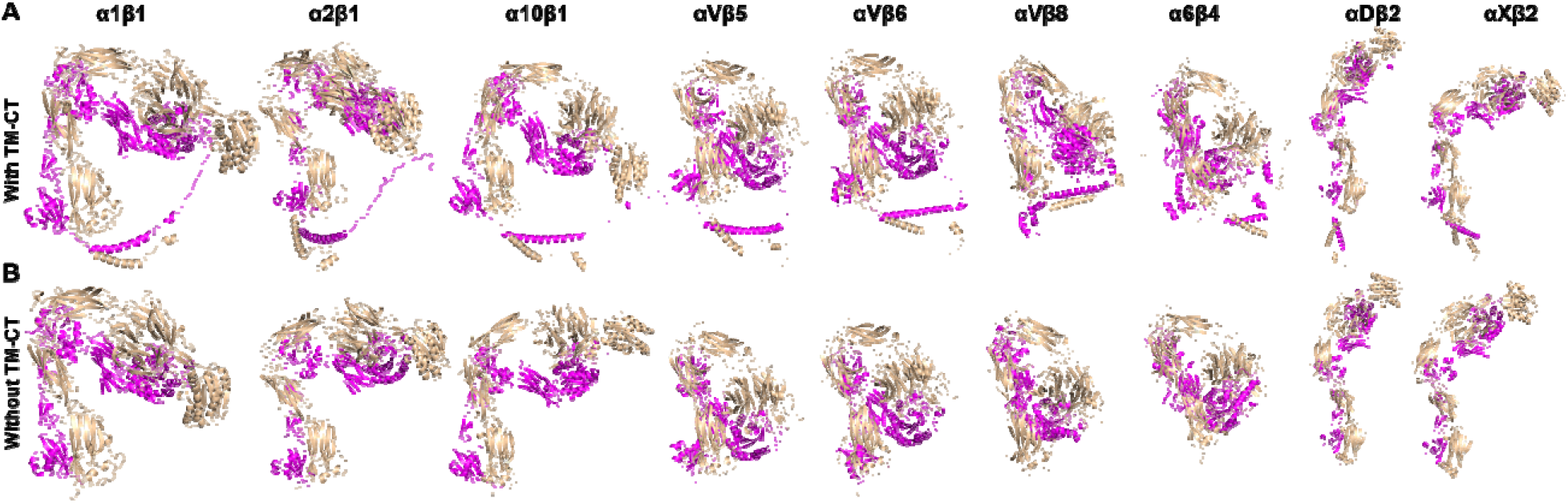
Comparison of structures predicted by AlphaFold2-multimer with and without transmembrane and cytoplasmic domains. (**A**) Full-length structures of certain integrins calculated by Alphafold2-multimer exhibit artificial contacts between the ectodomain and TM-CT domains. (**B**) The integrins shown in panel A were re-modeled by AlphaFold2-multimer in the absence of TM-CT domains. The structures were superimposed based on the calf-2 domain of the α subunit and are presented in the same orientation. The α and β subunits are shown in wheat and magenta, respectively.

We proceeded to investigate whether the provision of structure templates to AlphaFold2-multimer had any impact on the calculation of integrin structures. For this test, we selected α_5_β_1_, α_10_β_1_, α_V_β_8_, and α_X_β_2_ integrins. The template searching date was set to the year 2023, which allowed AlphaFold2 to use the integrin crystal structures as templates during structure calculation. The results show that all four integrin structures calculated by AlphaFold2-multimer, denoted as α_5_β_1_-2023, α_10_β_1_-2023, α_V_β_8_-2023, and α_X_β_2_-2023, exhibited a sharp bent conformation (**Fig. 8A-D**), closely resembling the crystal structure of bent α_IIb_β_3_ (**Fig. 8A**). For α_5_β_1_ integrin, this contrasts with the half-bent structure predicted by AlphaFold2-multimer with a search date set at the year 2000, denoted as α_5_β_1_-2000 (**Fig. 8A**). Indeed, the α_5_β_1_-2000 structure closely resembles the α_5_β_1_ cryo-EM structure (**Fig. 8A**). Similarly, the α_10_β_1_-2023 structure showed a bent conformation compared to the half bent α_10_β_1_-2000 structure (**Fig. 8B**). However, both α_V_β_8_-2023 and α_V_β_8_-2000 structures exhibited similar bent conformation (**Fig. 8C**). In contrast, the α_X_β_2_-2023 structure closely resembled the bent α_X_β_2_ crystal structure, while the α_X_β_2_-2000 structure appeared extended (**Fig. 8D**). These findings suggest that the inclusion of template structures can significantly influence the outcomes of structure prediction by AlphaFold2-multimer.

**Figure 8.**
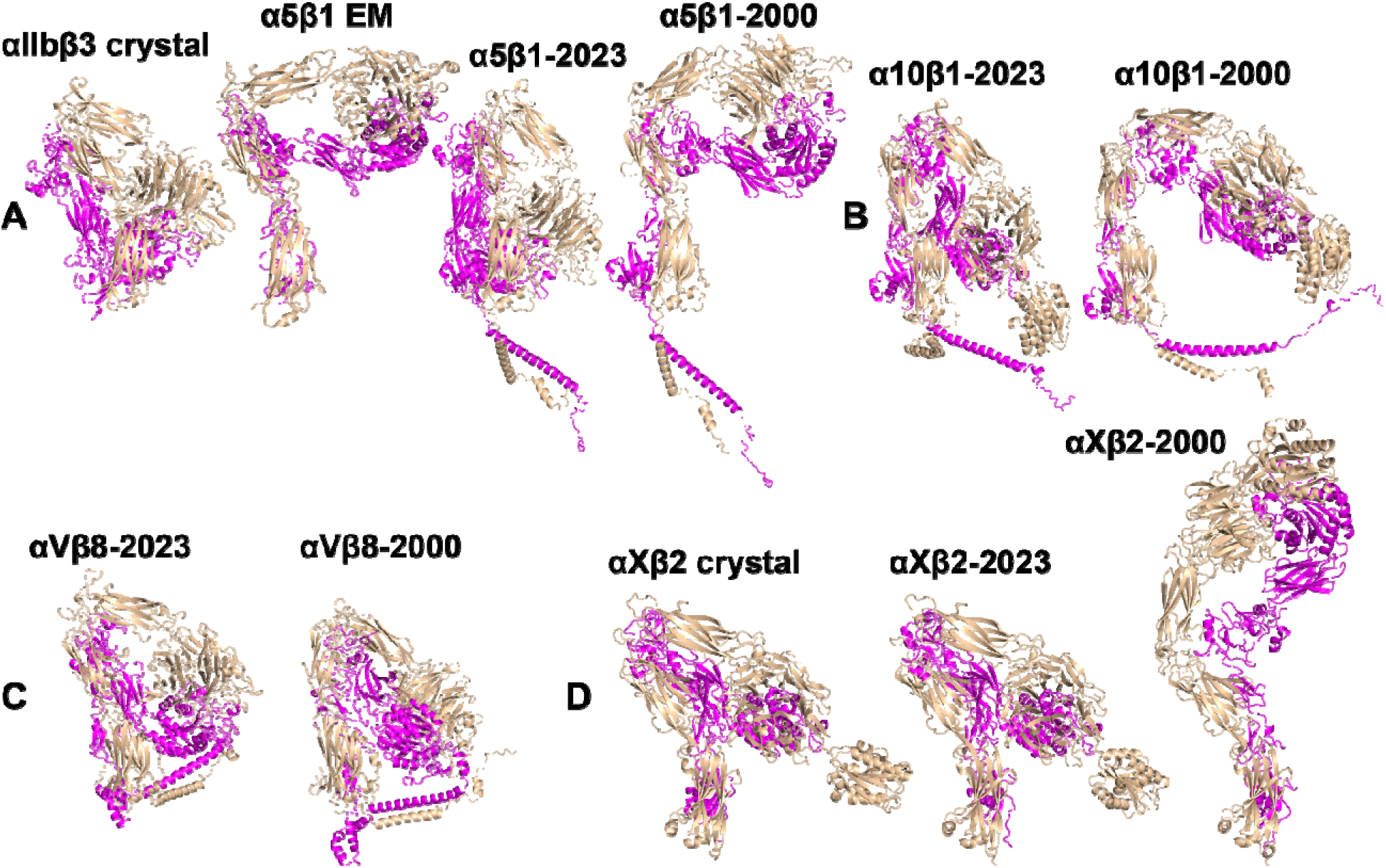
Comparison of selected structures predicted by AlphaFold2-multimer with and without enabling the option to use known structures as templates. (**A**) Full-length α_5_β_1_ structures were calculated by AlphaFold2-multimer with the max_template_date set to either year 2023 or 2000. Crystal structure of α_IIb_β_3_ (PDB 3FCS) and the cryo-EM structure of α_5_β_1_ (PDB 7NXD) were provided for comparison. (**B**) AlphaFold2 structures of full-length α_10_β_1_ with the max_template_date set to either year 2023 or 2000. (**C**) Full-length α_V_β_8_ structures modeled by by AlphaFold2-multimer with the max_template_date set to either year 2023 or 2000. (**D**) AlphaFold2 structures of ectodomain of α_X_β_2_. The crystal structure of α_X_β_2_ (PDB 4NEH) was shown for comparison.

### 3.5. Structures of integrin TM-CT domains

Despite the apparent simplicity of the sequences and structures of integrin TM and CT domains, experimental structure determination has been limited to the α_IIb_β_3_ TM-CT heterodimer. Through sequence alignment of the TM-CT domains from 8 β and 18 α human integrins, we identified conservative features at the TM, membrane-proximal (MP), and membrane-distal (MD) regions (**Fig. 9A**). These sequence conservative features were analyzed in the 24 integrin TM-CT structures calculated by AlphaFold2-multimer (**Fig. 9B**). Structure alignment revealed a high degree of structural similarity among the heterodimers at the TM domain, highlighting the conserved GXXXG motif in α and the conserved small G/A residue in β at the α/β TM interface (**Fig. 9B**). In the CT MP regions, the conserved GFFKR motif in the 18 α integrins all adopted a reverse turn conformation. The β CT MP regions, except for β_4_ and β_8_ integrins, displayed an α-helical structure extending from the TM α-helix, with the conserved Asp residue located at the α/β interface (**Fig. 9B**). The conserved β CT Asp residue is positioned proximal to the conserved Arg residue in the α GFFKR motif (**Fig. 9B**), which was proposed to form a salt bridge interaction [50]. The CT MD regions of both α and β subunits exhibited diverse disordered conformations, including the conserved NPXY motif responsible for binding talin (**Fig. 9B**). Despite not including any integrin TM-CT structure templates during the AlphaFold2-multimer calculation, the predicted α_IIb_β_3_ TM-CT structure closely resembled the experimentally determined structure (**Fig. 9C**). Additionally, we performed AlphaFold2-multimer modeling for the α_IIb_β_3_ TM-CT structure in the absence of the ectodomain, which showed a similar TM interface as the model generated along with the ectodomain (**Fig. 9D**). These results suggest that AlphaFold2-multimer is capable of accurately predicting integrin TM structures.

**Figure 9.**
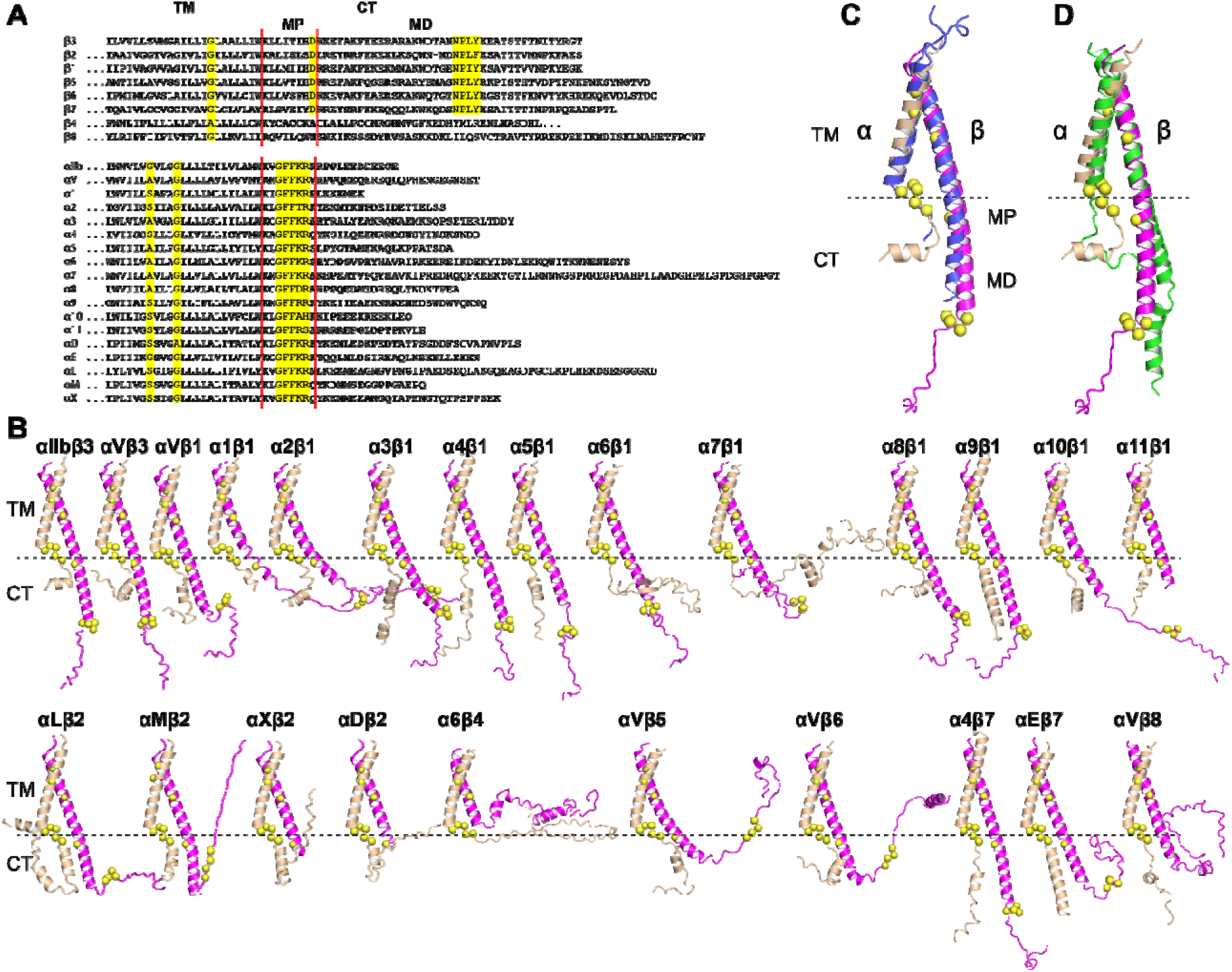
Structures of integrin transmembrane and cytoplasmic domains predicted by AlphaFold2-multimer. (**A**) Sequence alignment of human β and α integrin TM and CT domains. Conserved residues are highlighted in yellow. The boundaries of membrane proximal (MP) and membrane distal (MD) regions of CT domain are indicated. (**B**) AlphaFold2 structures of 24 human integrin TM-CT domains. The structures were superimposed onto the TM domain of α_IIb_ subunit and are presented in the same orientation. The conserved residues highlighted in panel A are depicted as yellow Cα spheres. (**C**) Superimposition of the AlphaFold2-predicted α_IIb_β_3_ TM-CT structure (in wheat and magenta) onto the heterodimeric structure of α_IIb_β_3_ TM-CT domains determined by disulfide crosslinking and Rosetta modeling (in blue). The α and β subunits are shown in wheat and magenta, respectively. (**D**) Superimposition of the α_IIb_β_3_ TM-CT structure predicted without (green) and with (wheat and magenta) the ectodomain.

## 4. Discussion

Our AlphaFold2-assisted structural analysis of the entire integrin family demonstrated the high performance of AlphaFold2 in predicting integrin structures. Sequence alignment analysis revealed an average sequence identity of 30-40% among the 8 β integrins and 20-40% among the 18 α integrins [38]. For individual integrin domains, such as the βI domain, the sequence identity can exceed 60% [10-12]. Since AlphaFold2 incorporates amino acid sequences, multiple sequence alignments, and homologous structures in its structure calculation [44], the high sequence identity among integrin domains likely contribute to the accurate prediction of integrin domain structures for most family members. This accuracy is exemplified by the striking similarity between the predicted structures and the experimental structures of α_IIb_β_3_, α_V_β_3_, and α_X_β_2_. Therefore, these predicted domain structures of integrins can be employed with a high level of confidence. They serve as valuable structural references for delineating domain boundaries when constructing integrin proteins for structural and functional studies. Moreover, the high-resolution structural models can be used for mutagenesis studies, identifying putative N-glycan sites and antibody epitopes, and interpreting data from functional experiments.

In addition to accurately predicting integrin domain structures, AlphaFold2 successfully modeled the global conformations of integrins in bent, intermediate (half-bent), or extended states. Remarkably, AlphaFold2’s calculations closely resembled the crystal structures for the bent conformations of α_IIb_, α_V_, α_X_, β_2_, and β_3_ integrins. According to the current integrin activation model, integrin activation involves a structural transition from a bent to an extended conformation. It is not readily known whether the intermediate and extended conformations predicted by AlphaFold2 for certain integrins represent a resting or activating state. However, these models are valuable for generating structure-based hypotheses that can be experimentally tested. For example, AlphaFold2 predicted an extended conformation for the single-chain α_10_ structure and a half-bent conformation for the α_10_β_1_ heterodimer. Consistent with these predictions, our flow cytometry assays, using mAb 9EG7 to report β_1_ integrin extension, suggest a constitutive extended conformation of α_10_ in the resting condition on cell surface, which was not previously expected. Notably, the single-chain α_7_ structure also exhibited an extended conformation, whereas the α_7_β_1_ heterodimer was modeled in a bent conformation. The 9EG7 binding assay indicated that α_7_β_1_ prefers a bent state under the resting condition on cell surface. However, the presence of activating Mn^2+^ induced more 9EG7 binding to α_7_β_1_ compared to other integrins such as α_3_β_1_ and α_9_β_1_ that were predicted to be bent by AlphFold2, suggesting that α_7_ integrin may be prone to becoming extended. These results imply that the overall integrin conformations predicted by AlphaFold2, especially for the underexplored integrins, can serve as reference structures for proposing functional assays.

AlphaFold2-multimer has demonstrated remarkable success in accurately predicting the structures of protein complexes, including those with transient interactions, multiple subunits, and large interfaces [51,52]. Here, we have shown that the AlphaFold2-multimer algorithm successfully predicts the structures of large complexes of integrin heterodimers. Furthermore, AlphaFold2 has exhibited excellent performance in predicting both fragment and full-length integrin structures. For instance, AlphaFold2-multimer’s modeling of integrin ectodomains remains consistent regardless of the presence or absence of TM-CT domains. Similarly, AlphaFold2-multimer effectively modeled the heterodimeric structures of integrin TM-CT domains, even in the absence of the ectodomain. Additionally, AlphaFold2 modeling may capture the conformational heterogeneity and intrinsically disordered regions, as observed in the predicted structures of integrin cytoplasmic domains.

Despite its relatively high accuracy, the integrin structures predicted by AlphaFold2 have apparent limitations. Notably, these predictions do not incorporate glycan structures and essential metal ions, both of which are vital components for integrin structure and function. Furthermore, the relative orientation between ectodomain and TM-CT domains cannot be correctly modeled by AlphaFold2. Additionally, our observations indicate that AlphaFold2-multimer predictions can be influenced by the homologous structures, potentially introducing bias when modeling the overall conformation of integrins. Therefore, we recommend conducting integrin structure calculations with AlphaFold2-multimer in the absence of template structures.

## 5. Conclusions

In summary, our family-wide structural analysis of integrins using AlphaFold2 showcases the remarkable capabilities of AlphaFold2 in modeling complex structures. The comprehensive structure database containing all 24 integrin heterodimers can be used as high-resolution structure resources for advancing both structural and functional studies within the integrin family.

## Author contributions

Heng Zhang: Investigation, Visualization, Writing-Reviewing and Editing.

Daniel S. Zhu: Visualization, Writing-Reviewing and Editing.

Jieqing Zhu: Conceptualization, Supervision, Visualization, Funding acquisition, Writing-Original Draft preparation.

## Acknowledgement

We thank the Research Computing Center at the Medical College of Wisconsin for providing the help and resources in running the AlphaFold2 program.

